# Traces of parenthood but not pregnancy loss in UK Biobank structural brain MRI data

**DOI:** 10.1101/2025.11.14.688235

**Authors:** Jonas L. Fischer, Alkistis Skalkidou, Birgit Derntl, Tobias Kaufmann

## Abstract

Pregnancy induces neuroanatomical changes in the human brain. Earlier studies detected traces of motherhood decades after childbirth. It is unclear whether these reflect persisting traces of pregnancy or effects of parenthood. We investigated effects of past birth and of pregnancy loss in women, and effects of fatherhood in men, using univariate and machine learning analyses on 205 regional brain volumes.

A group of mothers and an age-matched sample of nulliparous women (N=4357 per group, mean age 63 years) from the UK Biobank, with no past pregnancy losses, showed significant volumetric group differences in 14 regions at Bonferroni-adjusted α=.05. Likewise, we identified 18 significant group differences between age-matched samples of fathers and non-fathers of the same size (mean age 63.4), with 9 regions overlapping between sexes. Brain-wide association statistics for past live birth in mothers and those for fatherhood correlated (r = 0.55). XGBoost machine learning models trained to classify parenthood status separately in both datasets showed performance that was low, but significantly above chance (10-fold cross validation: AUC=0.56, p<1e-5 Motherhood classifier, AUC=0.54, p<1e-5, Fatherhood classifier, 10k permutations). We tested the motherhood classification model on an independent test sample comprising of four age-matched groups: 1. women who have never been pregnant, 2. women with past pregnancy loss but no live births, 3. women with live births but no pregnancy loss, and 4. women who experienced both. Class probability was significantly associated with live births, but not past loss.

These findings may suggest that neuroanatomical patterns of past childbirth partly also reflect traces of parenthood and not solely persisting traces of past pregnancy, although a more detailed characterization of pregnancy loss data would be needed for full confirmation of this interpretation. Therefore, further research is needed to quantify the extent and understand the nature of these changes, particularly considering the known vulnerability for mental disorders associated with reproductive events.

## Introduction

Pregnancy is a period of profound changes to the human body. These include structural and functional adaptations in the brain. Investigating neurobiological changes related to pregnancy is of high relevance due to the known vulnerability to mental disorders during this period^1,2^: Meta-analyses estimate antenatal depression to affect around 20% of pregnant women^3^, while the global prevalence of postpartum depression is estimated to be about 17% of mothers in the postpartum period^1^. Studies on mental health after pregnancy loss are scarce, however, one study found 36% of women to be highly symptomatic for depression, which was more than three times higher compared to pregnant controls^4^. The incidence of other mental disorders like anxiety, bipolar disorders or psychosis is elevated in the peripartum phase as well^2^. Establishing how disorders manifest in the brain during such a period of substantial neural restructuring^5,6^ may be key to a general understanding of the emergence of mental disorders as a multifactorial process^7^.

While the neurosciences are just beginning to characterize, quantify and understand pregnancy related changes in the brain, pre-to post-pregnancy grey matter volume reductions have been established and replicated^5,8–13^. An emerging hypothesis posits that these volumetric changes follow a U-shaped trajectory, where an initial decline in cortical grey matter during pregnancy is at least partially recovered in the postpartum period^8,10,14^. Affected brain regions include cortical regions like the prefrontal cortex as well as subcortical regions like the amygdala and hippocampus^5,8,10,15^. However, the actual extent and regional specificity of associations with pregnancy are highly dependent on the study, especially the time period investigated, with studies spanning different sections of the peripartum period from preconception to months or years postpartum^10,11,14,16–18^.

One possible major cause of brain adaptations are hormonal fluctuations. Pregnancy is known to cause substantial hormonal fluctuations in the human body, affecting also the brain^19^. It has been demonstrated that in murines, hormone-mediated neurobiological processes are directly linked with parenting behaviors^20,21^. Human studies found significant associations of specific sex hormones with grey matter volume trajectories^8,12^. However, the biological process of pregnancy is only one factor in this phase. Other factors include expectations of parenting, sociological adaptations as well as other changes to the environment of mothers. After birth, additional emotional adaptations are required, potentially causing cognitive challenges that could trigger structural and functional brain adaptations as well. Often, these experiences are shared between mothers that carry the pregnancy and their partners that may show similar effects of parenthood.

Studies on neurobiological adaptations in men transitioning to fatherhood have identified less pronounced, but still detectable cortical grey matter volume reductions either preconception or pre-birth to postpartum^22–26^. Interestingly, fathers have been shown to undergo transient hormonal shifts when their partners are pregnant^27^ but their correlation to brain volume trajectories is not established. Taken together, the literature indicates that not only pregnancy, but also parenting expectations or experiences can induce neural plasticity, potentially through different pathways. However, the only study to date investigating women that have never been pregnant, partnered with pregnant women, found a grey matter volume trajectory during their partners pregnancy that is comparable to nulliparous women^8^.

Still, one can hypothesize that parenting expectations and experiences influence brain volumes in both sexes, and the adaptations in mothers could be partly caused by pregnancy-related biological processes and partly by parenting expectations and experiences. It is established that specific brain networks are involved in parenting, and thus show increased activation in parents simulating caregiving in fMRI studies^28,29^. However, the extent to which parenthood as an environmental process or experience, apart from pregnancy, shape structural brain adaptations, is poorly understood. One way to study this is to investigate parents and non-parents later in life, when the biological processes around pregnancy and possibly breastfeeding are thought to have subsided.

A growing body of research suggests that associations of brain measures with parenthood exist until later in life, up to decades after the last birth^14,16,17,30–32^. It has been demonstrated that motherhood can be predicted from brain measures with a performance that is low, but significantly above chance level^16^. Studies are consistent in their findings that parenthood acts neuroprotectively. This is demonstrated, for example, by machine learning – based prediction of chronological age from the brain, resulting in a brain age biomarker^33,34^, which is consistently predicted to be younger for parents compared to non-parents^16,31^. The impact of parity on neuropathology is exemplarily shown by findings that multiparity is associated with decreased risk for Alzheimer’s disease^35^. Still, research on the trajectory of pregnancy-or parenthood related brain alterations later in life as well as their regional extent in the brain and potential consequences remains limited.

In the current study, we aimed to assess late-life effects of parenthood and pregnancy on brain structure as measured by T1-weighted MR imaging. We quantified parenthood effects on individual brain regional volumes, including cortical, subcortical, and ventricular features. Also, we aimed to investigate claims of late-life effects of pregnancy on the brain and tried to dissect whether these are actually associated with the biological process of pregnancy or rather caused by environmental factors like parenting experiences or other adaptations that come along with parenting.

To this end, we investigated effects of motherhood, fatherhood and pregnancy loss on 205 regional brain volumes in age-matched samples of parents and non-parents from the UK Biobank (UKB)^36,37^ in a cross-sectional manner. Using different comparisons of these effects, we aimed to distinguish pregnancy from parenthood effects. We deployed both univariate and multivariate streams of analysis to investigate possible patterns beyond single-region effects.

## Methods

### Data

This research has been conducted using the UK Biobank (UKB) Resource^36,37^ under Application Number 121699. We drew multiple samples from UKB, including all participants with available regional brain volume data, provided by UKB as imaging derived phenotypes (IDPs) from T1w brain MRI scans in the first imaging visit. The full documentation of data acquisition and initial processing is provided by UKB^38^. We split the participants according to sex (data field 31), which is taken from reports from the National Health Services from the UK and in some cases updated by participants. The terms “Female”, “Male”, “Women”, “Men”, “Mother”, and “Father” in the context of our work are inferred from this variable. We excluded data from all women who indicated unsure answers to the questions “Number of live births” (data field 2734) and “Ever had stillbirth, spontaneous miscarriage or termination” (2774). We also excluded participants reporting > 9 live births or > 11 total pregnancy losses based on extreme values in data distributions. We proceeded to split women into *mothers* and *non-mothers* based on their answer to “Number of live births”, leading to a binary variable “*motherhood”.* While women with no live births but pregnancy losses may generally also fall under classification of a mother, the term *motherhood* is used here for women that gave birth to a living child and are assumed to have undergone some form of a process of childcare, at least in most cases. *Pregnancy loss* was determined using data field 2774, from which we defined “*loss*” as “stillbirth, spontaneous miscarriage or termination”. The term “*loss*” therefore primarily indicates a prematurely lost pregnancy and does not necessarily indicate an event that is subjectively perceived as loss. The number of pregnancy losses was computed as the sum of answers to the data fields 3829, “Number of stillbirths”, 3839, “Number of spontaneous miscarriages” and 3849, “Number of pregnancy terminations”. Similarly, we excluded men with unsure answers to data field 2405, “Number of children fathered” as well as participants indicating > 9 fathered children, to gain a value range comparable to the dataset of women described above. Similar to women, we split men based on a binary “*fatherhood*” variable. From this data set, we generated an age-matched sample of mothers and nulliparous women with no reported pregnancy losses. We refer to this sample as “Female Training” as it was used to train machine learning models (see Table 1). Age-matching was performed based on integer age, according to data field 21003, “Age when attended assessment centre” (age bin: 1 year). Similarly, we created an age-matched sample of fathers and non-fathers (“Male Training”). Additionally, we generated another independent “Female Testing” dataset, which consisted of four groups:

1. women who have never been pregnant,
2. women with past pregnancy loss but no live births,
3. women with live births but no pregnancy loss, and
4. women who experienced both.

**Table 1:** Description of derived samples for analyses.

| Name | N | Age<br>years<br>(±<br>SD) | Description |
| --- | --- | --- | --- |
| Female<br>Training | 4357 x 2<br>groups | 63 ± 7.5 | Training Set for Women with no reported loss, age matched on binary live birth status (0-9 live births) |
| Female<br>Testing | 1297 x 4<br>groups | 61.9 ± 7.4 | Testing Set for Women, age matched on past loss and past live birth status (0-8 live births, 0-11 losses) |
| Male<br>Training | 4357 x 2<br>groups | 63.4 ± 7.7 | Training Set for Men, age matched on fatherhood status (0-9 children) |
| Female Full | 32765 | 64.4 ± 7.5 | Full Dataset of Women, not age-matched, with 0-9 live births and 0-11 total losses |
| Male Full | 29660 | 65.6 ± 7.7 | Full Dataset of Men, not age-matched, with 0-9 children fathered |

Table 1 summarizes main characteristics of the generated datasets. All samples were independent in the sense that each participant can only be included in one sample. While all groups in the individual samples were age-matched based on the procedure described above, the samples were not age-matched with the other samples. While the above-described samples were used for the main analysis given the clear assignment into age matched groups, we also performed a supplementary analysis using the full dataset available to us, with no age-matching applied and thus higher sample sizes (Table 1: Female Full and Male Full).

### Statistical analysis

For all above-described samples, we extracted regional brain volumes for 66 cortical brain regions defined by the Desikan-Killiany Atlas and 139 subcortical regions. Additionally, we used the computed estimated total intracranial volume (eTIV) as a covariate for the univariate analysis. A detailed description of these Imaging Derived Phenotypes provided by the UK Biobank can be found at https://biobank.ctsu.ox.ac.uk/crystal/crystal/docs/brain_mri.pdf.

For confound modelling in statistical association analyses, we used variables specifying the scanning site and an estimate of T1w scan based head motion of the participants. The full list of variables used, and their respective data field IDs can be found with the provided code of this analysis.

### Univariate analysis

To quantify parenthood effects on regional brain volumes, we computed linear regression models using the package *statsmodels* 0.14.4 for python 3.10.12. We defined regional brain volume *V(ROI)* as the dependent variable, and parenthood (binary yes/no), age, estimated total intracranial volume (eTIV), head motion and the assessment center site as independent variables. For all 205 regional brain volumes *V(ROI)*, we computed

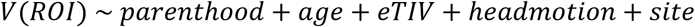

in both training datasets, separately. In our analysis, we focus on the association statistic *t* for the binary parenthood variable *t(parenthood)*, calculated as follows:

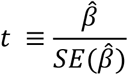

Thus, the *t* value quantifies how many standard errors the estimated coefficient differs from zero. To create a more intuitive understanding of the biological relevance of effects on a spectrum of neuroprotective to neurodegenerative, we flipped the sign of *t* values according to the direction of the effect of age on the respective brain volume for most of our subsequent analyses and figures. Thus, negative *t* values indicate an effect opposite to the effect of age that can be interpreted as potentially neuroprotective. We report brain regions for which the effect of parenthood surpasses significance thresholds, defined at α = 0.05, Bonferroni-adjusted for 205 tests. To test for the similarity of effects of parenthood in women and men, we computed the Pearson correlation of *t(motherhood)* and *t(fatherhood)* values across all brain regions.

Next, we preformed similar analysis using the Female Testing data, again computing linear regression models with the regional brain volume as dependent and either motherhood or past pregnancy loss as independent variable, with the same confounders as above. We repeated this 3 times>

1. Calculating *t(motherhood)* in a sample of women without past loss, using groups 1 (never pregnant) and 3 (live births, no loss) as introduced above (thus essentially comprising an independent replication of the analysis performed in the training data),
2. Calculating *t(loss)* in sample of women with no reported live births, using groups 1 and 2 (loss, but not live birth),
3. Calculating *t(loss)* in a sample of mothers (with reported live births), using groups 3 and 4 (both loss(es) and live birth(s)).

All three calculations were performed in samples with two groups of the same size and age-matched.

In addition, we performed a supplementary univariate analysis, using all data available to us, with no age-matching (Female Full and Male Full datasets). In the analysis of women, we modified the analysis to include *motherhood* and *loss* in the same regression model. In the analysis of men, we used binary *fatherhood* as dependent variable, similar as in the analysis in the Training data. The results of this analysis can be found in the Supplementary Material and Supplementary Figure 1.

Finally, to investigate the effects of parenthood in light of the effect of age on individual brain regions, we calculated and visualized the correlation of the *t* values of the predictors age and parenthood separately in both sexes.

### Machine learning analysis

Using the package xgboost 2.1.1 for python 3.10.12, we trained XGBoost classifier models to classify parenthood separately in both training datasets, maximizing the Area under the Receiver Operator Curve (ROC-AUC) as the training goal. We performed a parameter grid search for both models with the parameter grid shown in Supplementary Table 1.

We retained the models with hyperparameters leading to the highest mean ROC-AUC in a 10-fold cross-validation. We trained both final models on the whole training sample, yielding one *Motherhood Model* and one *Fatherhood Model*. To assess the generalizability of models across sexes we tested the performance of both trained models on the training data of the other sex. To test the probability of model performances occurring by chance, we performed permutation testing: We randomly permuted labels 10000 times and each time reran our cross-validation procedure.

We used the trained *Motherhood Model* to predict motherhood in an independent test sample consisting of the four groups of the Testing Sample (see Table 1). The predefined binary classification threshold is 0.5, meaning a prediction with a class probability *P(motherhood)* > 0.5 will be classified as a mother. Alongside binary prediction, we also investigated *P(motherhood)* as a continuous measure representing the estimated likelihood that a sample belongs to the class of mothers.

Using linear regression models, we tested for the association of class probability to the ground truth of *motherhood* and pregnancy *loss*. We included age, scanning site, and a measure of head motion, as confounding variables. Using the same confounding variables, we also tested for the association of class probability with number of live births, specifically in mothers, and number of pregnancy losses.

## Results

### Univariate analysis

For both sexes, the distribution of *t(parenthood)* values, quantifying the effect of parenthood on regional brain volumes as determined by linear regression (see Univariate Analysis), is generally shifted towards negative values (Figure 1A). This indicates parenthood tends to affect brain regions in the opposite direction to age. While the median of *t(fatherhood)* differs more strongly from zero than the median of *t(motherhood),* the most pronounced effects in mothers take more extreme values than those in fathers.

**Figure 1:**
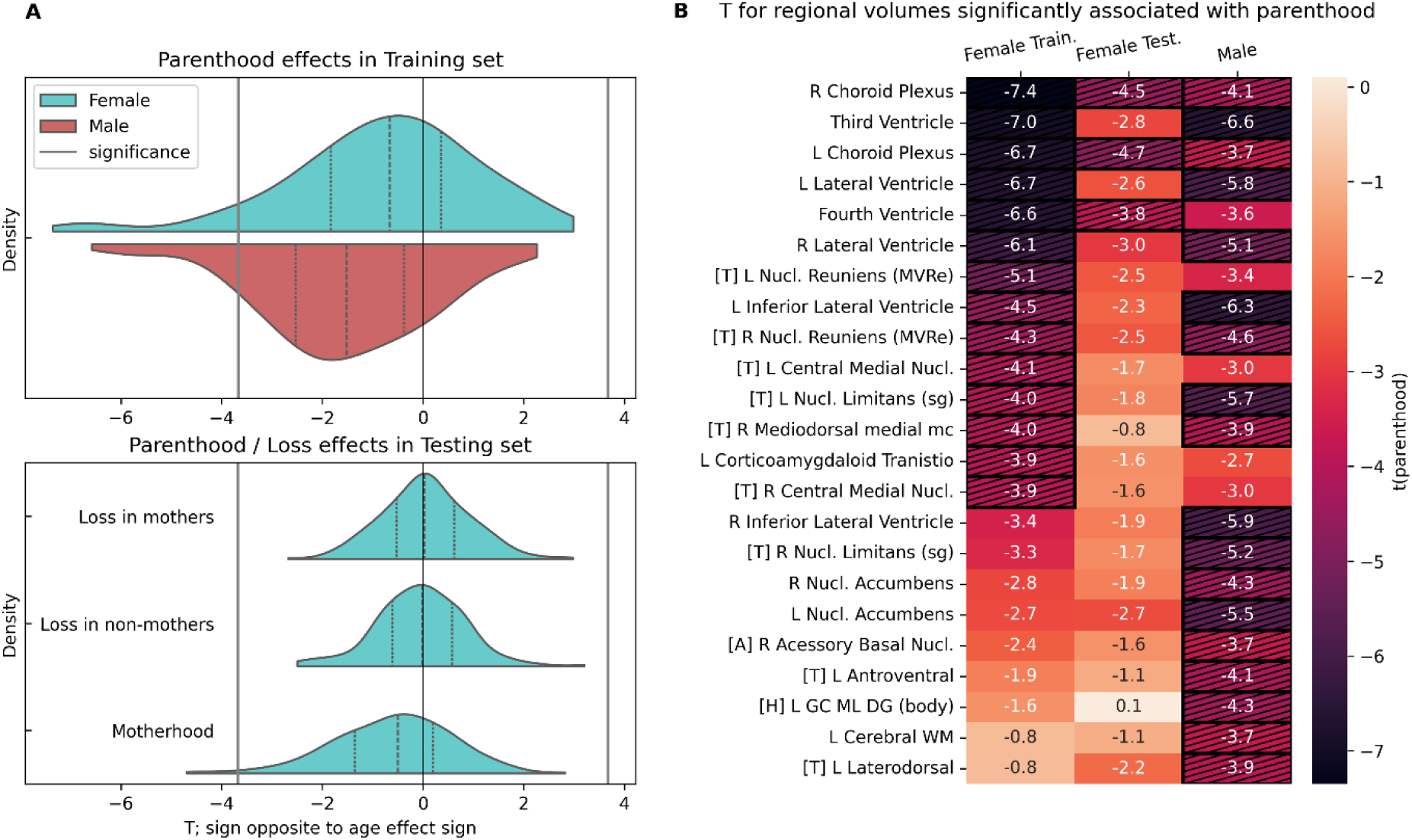
Univariate analysis reveals regional volume associations with parenthood in both sexes, but not with pregnancy loss. A: Distribution of t(parenthood) over 205 regional brain volumes separately for females and males (N = 8714 per sex) and distribution of t(loss) in mothers, t(loss) in non-mothers and t(parenthood), all in the testing sample (N = 2594 per model). B: T(parenthood) for females in the training and testing sample and for males, for regions that show a significant association with parenthood in either sex. Regions are sorted by t-values in the female training sample. All t values are modified according to the direction of the age effect (see Methods). Dashed cells indicate significance. Significance is defined using α = 0.05, Bonferroni-corrected for 205 tests. R = Right, L = Left, [T] = Thalamic Nucleus, [H] = Hippocampus, [A] = Amygdala.

We observed no visible effects when investigating the effects of pregnancy loss in the smaller Test Set. The distributions of t(loss) values in mothers and non-mothers are centered around zero (see Figure 1 A) and no values surpass significance level. However, in the same sample, we observe a distribution of t(motherhood) that is shifted towards negative values, replicating the results from the larger Training Set. In the independent Testing Set, the effect of motherhood surpasses significance in three brain regions, which are among the five regions with the strongest motherhood effects in the Training Sample.

In total, we identified 18 regional volumes significantly associated with fatherhood and 14 regional volumes significantly associated with motherhood. Of those, nine overlapped (see Figure 1 B). All of the significantly affected regional volumes were affected by parenthood in the opposite direction of age.

Many brain regions significantly associated with parenthood in either sex belong to either the ventricular system including the choroid plexi or are thalamic nuclei, as shown in Figure 1 B. Regional associations with motherhood discovered in the Training set mostly show the same effect directions in the smaller Testing set, except for the granule cell and molecular layer of the dentate gyrus (DG) in the body region (GC-ML-DG body) of the hippocampus, which had no notable motherhood association in the Testing Set.

In a supplementary analysis, we computed the effects of motherhood, fatherhood, and of pregnancy *loss* in larger datasets, omitting the age matching between groups to obtain higher statistical power. In these extended samples we reproduced the above-described findings with even more pronounced effects and with more values surpassing significance level compared to the main analysis. The distributions of *t(loss)* are centered close to 0, shifted to positive, with no values surpassing significance level (see Supplementary Figure 1). *T(motherhood)* and *t(fatherhood)* correlated with R ≈ 0.6, while *t(motherhood)* and *t(loss)* showed no notable correlation.

Figure 2 A shows that *t(motherhood)* and *t(fatherhood)* correlated with Pearson r ≈ 0.55 (*p =* 8e-18), indicating similarity of parenthood effects in women and men. Furthermore, we found that brain regional volumes strongly affected by age are also strongly associated with parenthood, indicating effects opposite to apparent aging in the whole brain rather than regional specificity (Figure 2B). Testing the correlation of parenthood and age effects, separately for both sexes, we observed strong correlations with Pearson r ≈ −0.8 (*p* = 2e-46 for women, *p* = 5e-44 for men).

**Figure 2:**
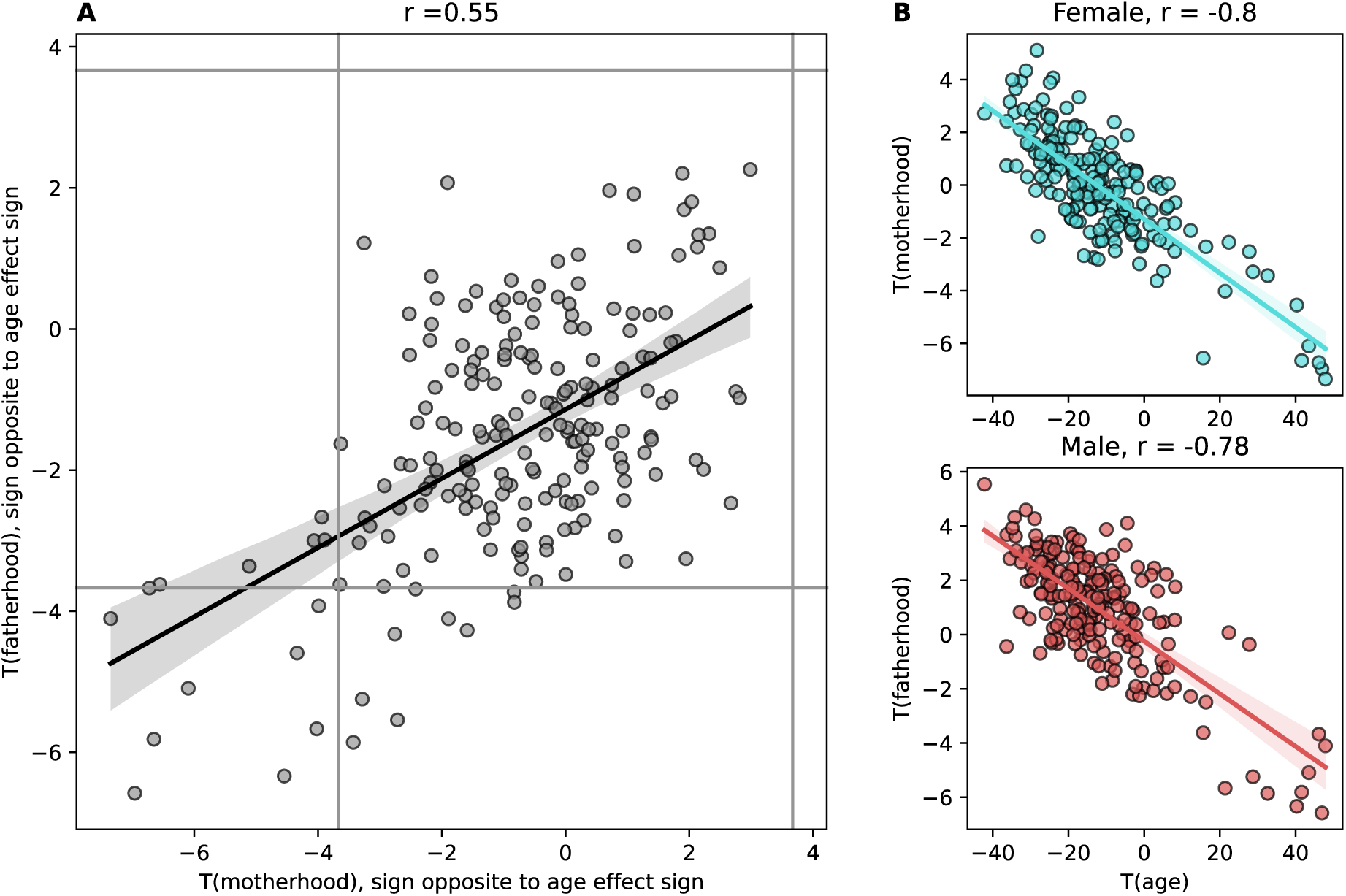
Effects of motherhood and fatherhood correlate with each other and with age. A: Scatterplot of t(motherhood) against t(fatherhood). Significance is defined using α = 0.05, Bonferroni-corrected for 205 tests. B: Scatterplot of t(parenthood) against t(age), by sex. Each dot represents one brain regional volume.

### Machine learning analysis

After optimizing hyperparameters (see Supplementary Material), our XGBoost-Classifier trained to predict motherhood (*Motherhood Model)* achieved a mean ROC-AUC of 0.56, while the *Fatherhood Model*’s mean ROC-AUC was 0.54, both measured by 10-fold cross validation in the training data. The *Motherhood Model* applied on an independent Testing set performed at an ROC-AUC of 0.56. Applying the models on data of the other sex yielded performance close to chance level, with ROC-AUC values of 0.5 – 0.51.

We found that both models performed significantly above chance level *(p <* 1 x 10^-5^), measured by randomly permuting training labels 10 000 times. The evaluation metrics for the final models, cross-sex performance, and permutation testing results, are shown in Figure 3.

**Figure 3:**
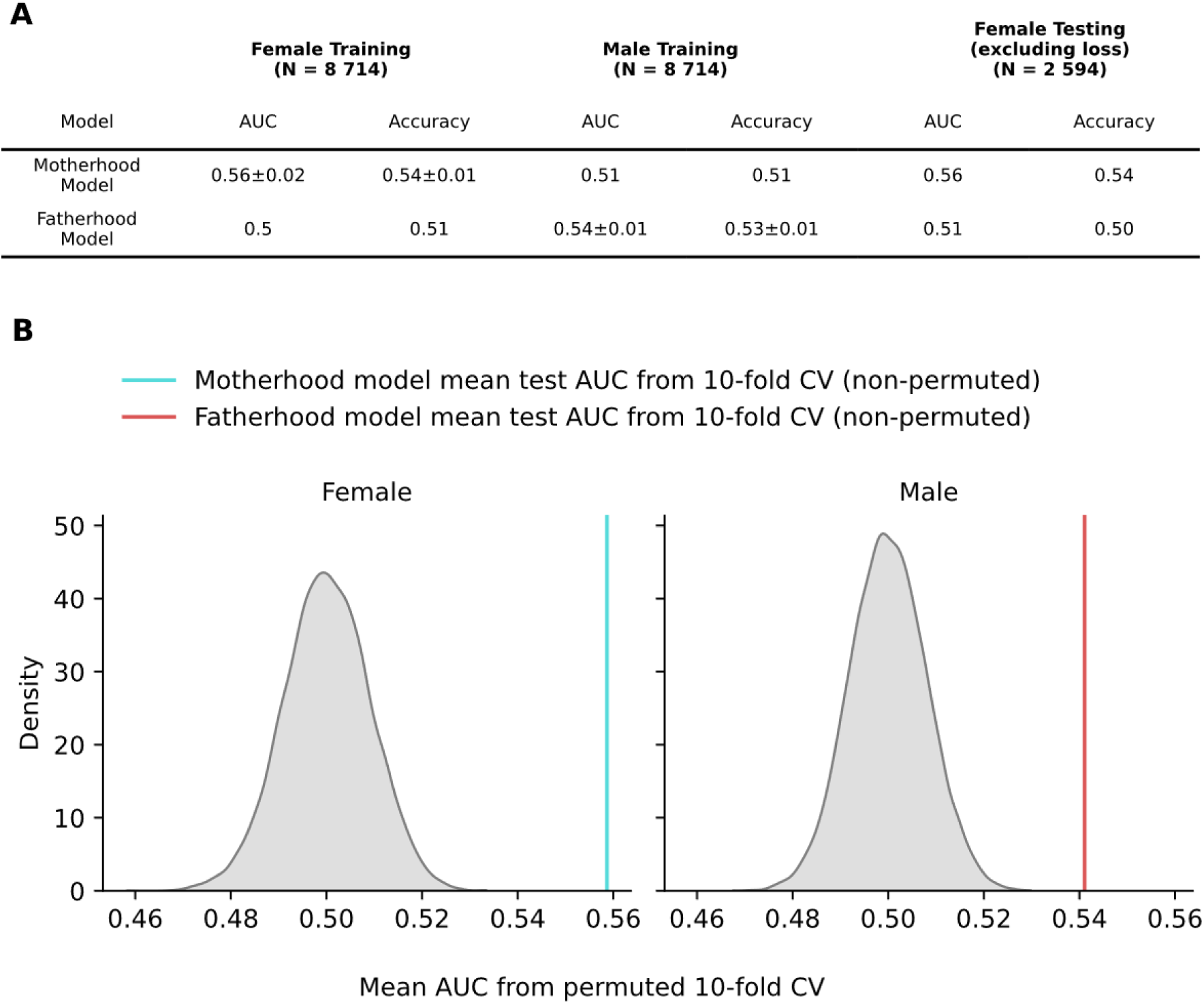
XGBoost classifier predicts parenthood with above-chance performance in both sexes. Table A shows ROC-AUC and accuracy for the models trained to predict motherhood respectively fatherhood. The performance metrics for the training data of the respective model is shown as mean + standard deviation from 10-fold cross validation. Figure B shows the distribution of mean ROC-AUC from model evaluation on the dataset with 10000 random permutations of features and labels. The vertical lines display mean model performance on original training data.

Investigating the degree to which past loss is associated with multivariate patterns of motherhood, we applied the model to the independent Testing Sample also containing women after pregnancy loss. The distribution of class probabilities for the four different groups shows high visual similarity (Figure 4A). While the distribution for the two groups of women without live births looks almost identical, the distribution of women who reported live births, but no pregnancy *loss*, is shifted towards positive values compared to the two groups with no live births. The distribution for group 4, women who reported both live births as well as pregnancy losses, is shifted slightly towards negative compared to group 3, who did not report pregnancy losses.

**Figure 4:**
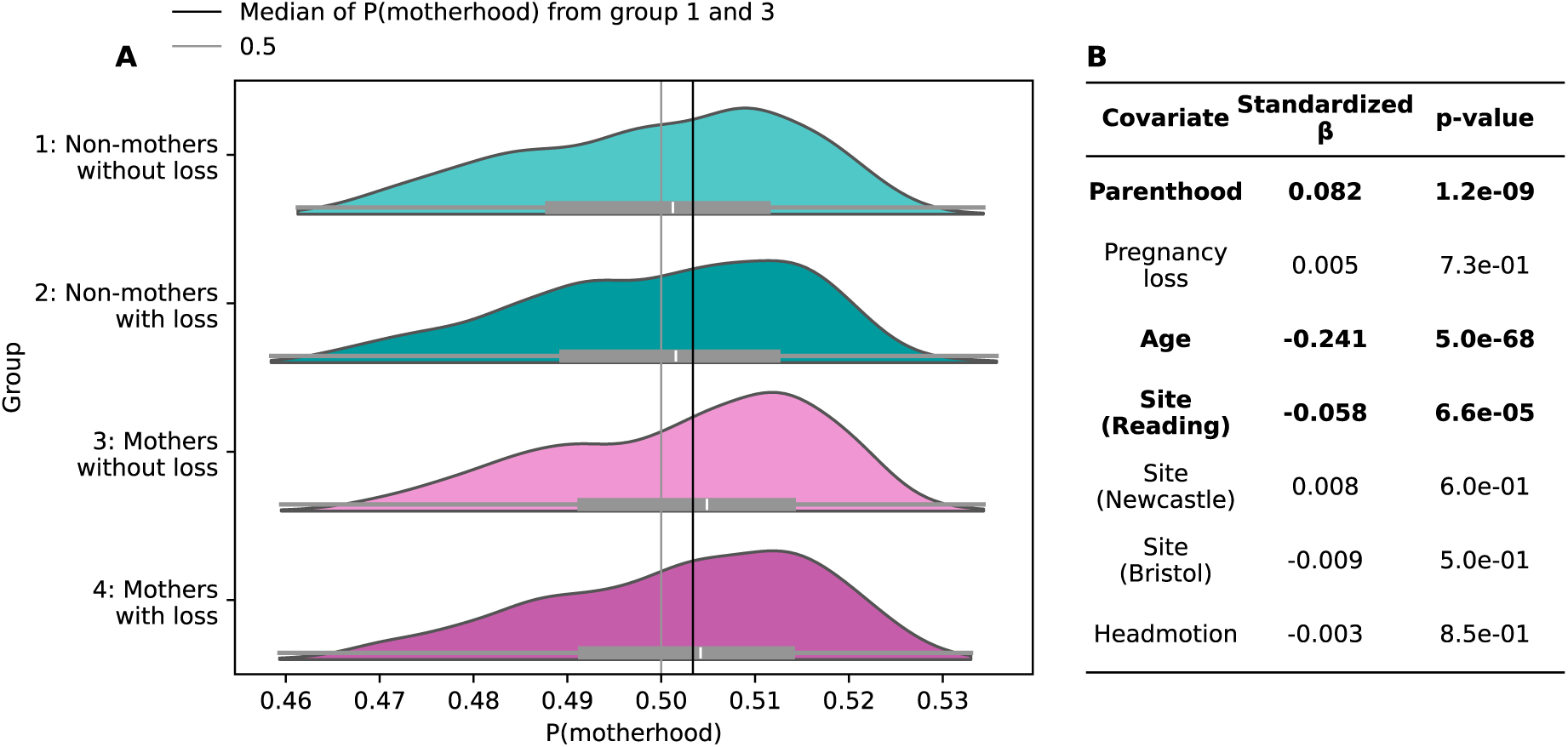
Model predicted class probability P(motherhood) is significantly associated with parenthood status, age and site, but not pregnancy loss. A: Distribution of model predicted class probability in the testing set, separately for the 4 age-matched groups. B: Covariates, coefficients, and p-values for the linear regression model with class probability as the independent variable (N = 5188). Bold numbers denote significant independent variables (p < 0.001).

Importantly, while class probabilities were associated with parenthood, they were not significantly associated with a past pregnancy loss. Class probabilities were positively associated with motherhood (*p* = 1e-9, see Figure 4B), as expected for a model classifying motherhood. We computed these associations with linear regression models using age, site, and a measure of head motion as covariates. Age was negatively associated with class probability, as well as the scanning site of Reading, indicating lower class probabilities assigned to volumes obtained at this scanning site. The full regression results are shown in Figure 4B.

Younger women tended to be classified with a higher probability of being mothers (Figure 5 A). The fitted regression slopes appear parallel for mothers and non-mothers, indicating no important interaction effect between age and actual motherhood. In further linear regression analyses, we found that class probability is significantly associated with number of live births, also when considering only actual mothers. As shown in Figure 5B, class probability tended to be higher for multiparous women, compared to women with one to two live births. Linear regression results showed that the association of class probability with number of live births was significant (*p* = 1e-10) also when considering only the subgroup of mothers (*p* = 0.02), while there was no significant association between class probability and number of pregnancy losses (*p =* 0.24).

**Figure 5:**
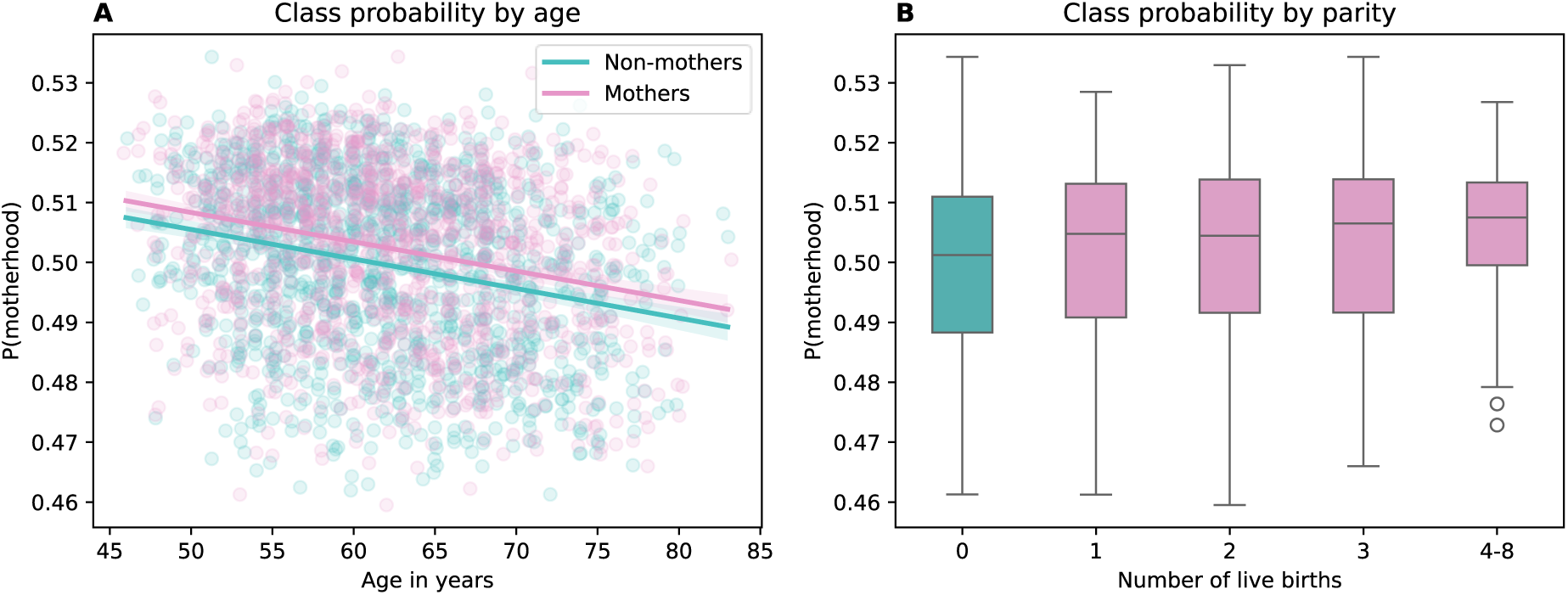
Class probability is higher for younger women and multiparous mothers. A: Linear model fit with class probability as dependent and Age in years as independent variable, separately for mothers and non-mothers. Lighter bands correspond to the 95% confidence interval. B: Boxplots of the distributions of class probability for number of children. Women with 4 to 8 children were grouped to obtain meaningful sample sizes. In both visualizations, women with reported pregnancy loss were excluded.

## Discussion

### Univariate analysis

We computed effects of parenthood on 205 individual brain regional volumes in an age-matched UKB dataset of both sexes. Parenthood effects on brain regional volumes show a strong negative correlation with age effects, potentially indicating a neuroprotective effect of parenthood. This is consistent with previous findings of younger brain age in parents^30,31^ and parity effects that are protective of brain function^32^.

With a rather conservative multiple testing correction defined by Bonferroni, we here identified 14 regions significantly associated with parenthood in women and 18 in men, of which nine overlapped. Brain-wide association statistics of fatherhood and motherhood correlated between sexes, indicating a level of similarity of these parenthood effects in mothers and fathers.

Most of the regions affected in both sexes were regions of the ventricular system including the choroid plexi, which produce cerebrospinal fluid (CSF) from the blood^39^. The FreeSurfer-based choroid plexus segmentation has been described as being less accurate compared to other methods^40^, warranting follow up studies with more sophisticated segmentation methods. Ventricular enlargement is associated with neurogenerative diseases^41^ as well as lower cognitive function in healthy individuals^42^. The volume of the fourth ventricle has been found to be a significant feature in age prediction from the brain in UKB data^33^, highlighting its association with age-related brain trajectories. Other regions significantly associated with motherhood include multiple thalamic nuclei. The thalamus plays a major role in relaying sensory information and processing motor information^43^. De Lange et al. have found a highly significant association of motherhood on a cluster of mostly limbic regions^30^, also including the thalamus. Fatherhood was significantly associated with total cerebral white matter and the nucleus accumbens volume, which is part of the reward system^44^. Other regions of the reward system show increased activation in fathers listening to their infant’s cries^45^ and in mothers with infant stimuli^46^ suggesting a potential role of these structures in reacting to offspring cues. Taken together, the prominent associations in ventricular and choroid plexus regions may indicate that parenthood exerts diffuse, whole-brain effects that are most readily detected in structures sensitive to global brain aging^47^. This interpretation is consistent with the idea that these findings reflect cumulative rather than highly localized effects.

Motherhood showed higher effect sizes on the regional volumes with most pronounced associations compared to fatherhood. Yet, the total number of associated regions was higher for fathers than mothers. These results were unexpected, as mothers in the past and present have been shown to devote significantly more time to childcare compared to fathers^48^ and undergo the biological process of pregnancy, introducing additional long-lasting brain adaptations^5,11^. Indeed, our results contradict another study investigating parenthood associations in grey matter in data obtained from the UK Biobank, reporting regional grey matter volume differences in mothers compared to non-mothers, but not in fathers compared to non-fathers^14^. However, the study used a different brain segmentation atlas from the one used here, and the analysis focused on specific regions of interest previously identified in studies investigating women during the peripartum phase. Interestingly, another study investigating parenthood effects on brain function in UKB functional MRI data, found numerically stronger parenthood effects in men compared to women^32^. The authors discuss that this result may reflect either biological differences, such as more general sex differences in the brain^19,49^, specific biological differences in this sample (e.g. the impact of pregnancy) or gender- and sex-driven differences in parenting and caregiving expectations.

We did not identify any effects of past pregnancy loss. We could however replicate motherhood effects in an independent dataset of the same size as the one used for testing for pregnancy loss effects. Thus, there were no lasting neuroanatomical traces of pregnancies that ended prematurely in individual brain regional volumes. Other studies on UK Biobank data have not systematically described how they analyzed pregnancy loss, but also report not finding significant effects^14^. To describe brain volume trajectories after pregnancy loss, studies with a defined and accurate characterization of pregnancy losses, investigating longitudinally and starting shortly after the loss, would be needed.

We employed a rigorous age-matching procedure to our data to minimize the risk of conflating age and parenthood related effects, at the cost of substantially reducing our sample size. We matched women and men using a 1-year age range on the binary *parenthood* variable for the Training Datasets and women on the *parenthood* and *loss* variables for the Testing Set. Using this procedure, we could identify significant motherhood and fatherhood effects on brain regional volumes, but not pregnancy loss effects. We observe the same results, but more pronounced, when using the full datasets without the age-matching procedure, potentially indicating that we could observe even stronger effects with more statistical power.

### Machine learning analysis

Our multivariate classifier trained to predict motherhood from regional brain volumes performed with low accuracy, but significantly above chance. Our classifier performed in line with, or better, than classifiers trained for the same task in similar data in other studies^16,31^. We were thus able to reproduce previous results, including data from a new UKB data release. Our classifier trained to predict parenthood in women does not perform above chance level in men and vice versa. This implies that while the univariate effects of parenthood show correlation between sexes, the multivariate patterns learnt by the classifier do not generalize to the other sex.

We applied our classifier trained to predict motherhood on an independent test sample, containing, next to nulliparous women and mothers without pregnancy losses, also mothers with and without pregnancy losses. We found that the model’s predicted class probability was associated with motherhood, but not pregnancy loss. While we cannot rule out the presence of multivariate patterns of past pregnancy loss, we can conclude that if they are present, they do not take the same form as motherhood patterns learnt by the classifier.

Our results indicate that late-life effects of parenthood reflect traces of the experience, challenge and task of motherhood or the adaptations that come with it rather than solely traces of the biological process of pregnancy. This is supported by the fact that both motherhood as well as fatherhood are associated with persisting volumetric effects that are correlated across sexes. Pregnancy loss is neither associated with differences in individual brain regional volumes nor a multivariate class probability of motherhood.

### Limitations

Due to the cross-sectional nature of the study, the results are to be cautiously interpreted. We cannot rule out that the identified effects of parenthood represent factors that influence the probability of becoming a parent itself. This selection bias could be direct, i.e. men and women more prone to have a partner and become parents already show distinct differences in the brain. It is also possible that there are other factors, for example environmental influences like socioeconomic status or lifestyle, or biological/genetic factors that affect the probability of becoming a parent and also shape differences in regional brain volumes. Furthermore, the identified effects could also be caused by lifestyle changes that come with parenting, for example an adoption of a healthier – and neuroprotective – diet.

The items we made use of to define pregnancy loss do not capture all potentially relevant information on this topic and are based on self-reports. By classifying early loss, late miscarriages, and abortions as pregnancy *loss*, we grouped potentially heterogenous events based on the fact that pregnancy was ended prematurely. Information that would be very relevant in the context of this analysis include the gestational age at which the loss occurred. Depending on that, the biological adaptations might not reach the same progress as those in women undergoing pregnancy ending in live births. While an analysis comparing effects of parenthood with effects of only stillbirths would be beneficial in this study, we decided not to pursue this due to the small number of participants reporting stillbirths, but no live births (n = 38). Due to the prevalences of pregnancy losses at different time points, we have to assume that the majority of pregnancy loss investigated here are short pregnancies. Thus, it is very much possible that the observed null-effects of pregnancy loss are due to a large part of pregnancies being not developed enough for major structural changes to take place.

Also, the individual experiences of these events can differ drastically. Additional items on the subjective experience of these events would be beneficial. For example, a strong grieving process^4,50,51^ could potentially induce lasting neuroanatomical effects, masking or acting against pregnancy-related effects. Men answering the UKB questionnaires did not answer questions about pregnancy losses of their partners at all. It is possible that some would include lost pregnancies in their answer to the question how many children they fathered. Additional and accurate questions about reproductive history would also be very beneficial for our analysis of fatherhood in men.

The UK Biobank resource covers a large population, however, is mostly comprised of White Europeans, thus, we cannot be sure that these findings generalize to other populations. Additional information from questionnaires that would be valuable for our analysis would be, for example, whether some participants also became grandparents, as this might influence possible brain changes caused by sociological adaptations further. More questionnaire data on same-sex couples or self-reported gender would also be beneficial. The nature of human parenting is highly diverse and parenting situations may be highly individual and dependent on sociocultural settings, socioeconomic structures or individual family structure and support network composition. These individual situations can be hypothesized to mediate the relationship between parenting and brain adaptions. The population and the data collected do not fully capture this diversity.

## Conclusion

In conclusion, our study reproduced previous results of a potentially neuroprotective effect of motherhood^16,30–32,35^ and extends it also to fatherhood. Also, we were able to reproduce results of parenthood being classifiable from brain regional volumes at levels close to, but above chance in late life^16,30^, including participants from a new data release compared to previous publications. Using comparisons of motherhood, fatherhood, and pregnancy loss effects on the brain, we provide evidence that brain differences between mothers and non-mothers decades after childbirth might be associated with the process of parenting as a change to the individual’s environment rather than the biological process of pregnancy. To further disentangle these effects, longitudinal studies of populations that undergo either pregnancy or motherhood – like women with lost pregnancies or foster resp. adoptive parents – will be of high relevance.

## Acknowledgements

Kaufmann received support from the European Research Council (ERC CoG, #101086793, HealthyMom), the Research Council of Norway (#323961) and the German Research Foundation (IRTG 2804). He is a member of the Machine Learning Cluster of Excellence, EXC number 2064/1, Project number 39072764. Brain imaging data was processed on the BMBF-funded de.NBI Cloud as part of the German Network for Bioinformatics Infrastructure (031A537B, 031A533A, 031A538A, 031A533B, 031A535A, 031A537C, 031A534A, 031A532B). This research has been conducted using the UK Biobank Resource under Application Number 121699. The authors thank UK Biobank participants for making their data available to the research community.

## Data and code availability

This research has been conducted using the UK Biobank Resource under Application Number 121699. The data is available upon request to the UK Biobank.

All code that was used for the analyses is will be available on GitHub upon acceptance of this manuscript (github.com/jonlfischer).

## Author contributions

Jonas L. Fischer: Conceptualization, Formal analysis, Data Curation, Visualization, Writing—Original Draft, Writing - Review and Editing. Alkistis Skalkidou: Supervision, Writing—Review and Editing. Birgit Derntl: Supervision, Writing—Review and Editing. Tobias Kaufmann: Conceptualization, Data Curation, Funding acquisition, Project administration, Resources, Supervision, Writing—Original Draft, Writing—Review and Editing.

