## Supplementary Materials for "Traces of parenthood but not pregnancy loss in UK Biobank structural brain MRI data"

### Supplemental material

**Univariate analysis**

We performed an additional analysis to make use of our full dataset without our rigorous age-matching procedure. In our Full Dataset of men (see Table 1), we computed the effects of *fatherhood* using the same linear regression model as described above (see Methods). Similarly, we computed the effects of *motherhood* and pregnancy *loss,* however, in the same linear model, thus calculating

$$V(ROI) \sim motherhood + loss + age + eTIV + headmotion + site$$

The resulting distributions of *t(motherhood), t(fatherhood)* and *t(loss)* are shown in Supplemental Figure 1.

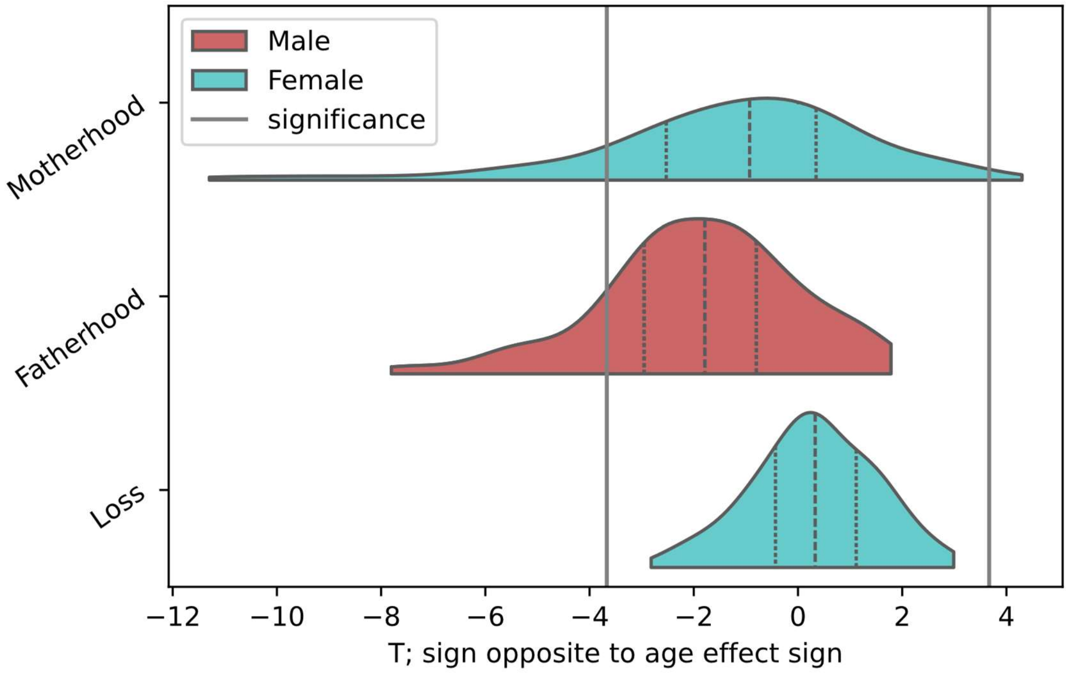


*Supplementary Figure 1: Distributions of t(motherhood), t(fatherhood) and t(loss) from supplementary analysis, omitting age matching. T values result from linear regression models in the Female Full and Male Full dataset.*

Similarly to our main analysis, we find negatively shifted distributions of *t* values for motherhood and fatherhood and numerous brain regions surpassing significance level. This pattern is more pronounced compared to our main analysis. Different from our main analysis, there are brain regions where the motherhood effect surpasses significance where the motherhood effect acts in the same direction of age. The distribution of *t(loss)* is shifted towards positive values, underscoring no neuroprotective effects of lost pregnancies could be identified. In this analysis, *t(motherhood)* and *t(fatherhood)* correlate with Pearson's r = 0.6.

#### Machine learning analysis

In our machine learning analysis, we performed a parameter grid search, building models with different variations of hyperparameters for the XGBoost classifier model. We then measured performance by 10-fold cross validation on the training set. The tested hyperparameter grid and the final values resulting in the best performance are shown in Supplemental Table 1.

| Parameter | Values in Grid Search | Value in final Motherhood Model | Value in final Fatherhood Model |
|---|---|---|---|
| eta | 0.001, 0.005, 0.01, 0.05, 0.1 | 0.001 | 0.05 |
| n_estimators | 25, 50, 100, 200, 400 | 400 | 50 |
| subsample | 0.6, 0.8, 1.0 | 0.6 | 0.6 |
| colsample_bytree | 0.6, 0.8, 1.0 | 0.6 | 0.6 |
| max_depth | 2, 3, 5 | 5 | 3 |
| min_child_weight | 0.5, 1, 3 | 3 | 3 |

*Supplementary Table 1: XGBoost parenthood classifier hyperparameters tested in grid search, and final hyperparameters for both models, selected based on highest ROC-AUC.*
